# Anterior cingulate cortex hypofunction causes anti-social aggression in mice

**DOI:** 10.1101/2020.10.07.328765

**Authors:** S. van Heukelum, K. Tulva, F. Geers, S. van Dulm, I. H. Ruisch, J. Mill, J. F. Viana, C. F. Beckmann, J. K. Buitelaar, G. Poelmans, J. C. Glennon, B. A. Vogt, M. N. Havenith, A. S. C. França

## Abstract

Controlling aggression is a crucial skill in social species like rodents and humans, and has been associated with anterior cingulate cortex (ACC). Here, we demonstrate a causal link between ACC hypofunction and failed aggression control in BALB/cJ mice. We first show that ACC in BALB/cJ mice is structurally degraded: Neuron density is decreased, with pervasive neuron death and neuro-toxic astroglia. Gene-set enrichment analysis suggested that this process is driven by neuronal degeneration, which then causes toxic astrogliosis. cFos expression across ACC indicated functional consequences: During aggressive encounters, ACC was engaged in control mice, but not BALB/cJ mice. Chemogenetically activating ACC during aggressive encounters drastically suppressed anti-social aggression but left adaptive aggression intact. The network effects of our chemogenetic perturbation suggest that this behavioural rescue is mediated by suppression of amygdala and hypothalamus and activation of mediodorsal thalamus. Together, these findings highlight the causal role of ACC in curbing anti-social aggression.

## Introduction

Aggression is an evolutionarily conserved behaviour present in most species including humans. What’s more, when applied in a context-appropriate way, e.g. by defending against intruders, aggression is a crucial survival skill [1]. In contrast, dysfunctional aggression such as abusive and anti-social behaviour leads to severe long-term personal and societal harm [1, 2], and is often observed in the context of different psychiatric disorders, including borderline personality disorder, psychopathy and conduct disorder [3]. What could be the neuronal mechanisms underlying maladaptive aggression?

Aggressive behaviour can generally be thought of as the product of two interconnected and complementary – even at times antagonistic – cortical and subcortical networks. One network, also referred to as the threat circuit [4], centres around the bed nucleus of the stria terminalis as well as the (ventromedial) hypothalamus and amygdala, which integrate information on potential threats, and initiate aggressive output via the periaqueductal grey (PAG) [4, 5]. In line with this, studies in rodents show that stimulating the hypothalamus [6–8] and specific neural populations within the amygdala [9, 10] triggers aggressive behaviour. A second network including cingulate cortex, lateral septum and insula is thought to mediate the top-down regulation of aggression [11–13]. Within this network, cingulate cortex, and specifically anterior cingulate cortex (ACC), seems to have a crucial gating function for aggressive behaviour: In wild-type rodents, inhibiting ACC leads to sudden bursts of aggression during social encounters [14].

Given that maladaptive aggression can mainly be thought of as aggression applied in the wrong context and/or not proportionally to the situation, it seems plausible that regulatory areas like ACC play a key role both in the development and treatment of dysfunctional aggression [2]. This notion is supported by several correlative observations. For instance, studies in patients with pathological aggression demonstrate alterations in ACC volume [15, 16] as well as ACC activity [16, 17]. Similarly, studies of ACC structure in a mouse model of maladaptive aggression, the BALB/cJ strain, show a volumetric increase of ACC compared to the non-aggressive BALB/cByJ control strain [18], as well as a decrease of Parvalbumin- and Somatostatin-expressing interneurons across cingulate cortex [19].

Despite these indications that dysfunctional aggression correlates with altered ACC structure and function, there has been surprisingly little work directly testing the causal role of ACC in dysfunctional aggression. Here, we provide this causal link by showing that ACC activation directly curtails anti-social - but not socially expected - aggression in BALB/cJ mice. We first characterized structural and functional alterations of ACC in the BALB/cJ strain, and discovered pervasive neuron death, accompanied by toxic glial processes, which resulted in stark reductions of neuronal density compared to control mice. Gene-set enrichment analysis of genetic differences between BALB/cJ and control mice supported these data by highlighting genes involved in neuronal degeneration. On a functional level, cfos expression indicated that BALB/cJ mice failed to activate ACC during aggressive encounters. Most importantly, we found that chemogenetically activating ACC in BALB/cJ mice was enough to prevent antisocial aggression, while having a smaller effect on socially expected aggressive behaviours. Analysis of cfos expression in downstream areas indicated that this behavioural rescue was most likely due to decreased activity within the amygdala and hypothalamus and increased activity in mediodorsal thalamus, suggesting a re-instatement of top-down modulatory control on the threat circuit. To our knowledge, these results for the first time establish a direct causal link from neuronal activity levels in ACC to the successful control of maladaptive aggression.

## Results

### Structural degradation in ACC of aggressive BALB/cJ mice

In line with previous reports [18–20], BALB/cJ mice consistently behaved more aggressively than control BALB/cByJ mice in the resident-intruder (RI) test – a paradigm in which an unfamiliar ‘intruder’ mouse is introduced into the home cage of another mouse (the resident) for 5-10 minutes per day, with a new encounter taking place daily for five days in a row (see Supp. Mat. Fig. 1). In this scenario, BALB/cJ mice not only attacked faster and more frequently than control mice, they also targeted vulnerable body parts like the face, abdomen and neck - a behaviour that is not expected during territorial conflicts in mice and can therefore be considered as anti-social in this context [18, 19]. To exclude possible litter effects, we tested several independent cohorts. We also allowed a sub-group of mice (n = 5 per strain) longer interaction times to exclude the possibility that control mice show similarly aggressive behaviour, but with a longer attack latency. As Supplementary Figure 1a shows, BALB/cJ mice behaved more aggressively under all circumstances.

Based on the hypothesis that excessive anti-social aggression might be due to a failure of the control circuits, we examined structural changes in ACC. We first performed a Cresyl-violet staining which enabled us to assess structural features of neurons (Fig. 1a). Across all layers of ACC, we found high rates of pyknotic neurons (Fig. 1b, repeated measure ANOVA, with strain as between-subject factor and layer as within-subject factor: *F*(1, 31) = 18.69, *p* < .001, *η2* = .38; post-hoc t-tests per layer: all p < .01). During pyknosis, the chromatin in the nucleus of a cell condenses, an irreversible process that occurs during cell death [21]. As such, our data show that a large number of neurons are dying within ACC of BALB/cJ mice. In line with this, unbiased stereological counting with the optical fractionator tool of the program StereoInvestigator (MBF Biosciences) revealed that neuron densities in ACC were drastically decreased across all cortical layers (Fig. 1c, repeated measures ANOVA: F(1, 31) = 23.05, p < .001, η2 = .43; post-hoc t-tests per layer: all p < .01). In BALB/cByJ mice, ACC contained on average approximately 35% more neurons (3246 neurons/mm2) than in BALB/cJ mice (Fig. 1C). To exclude the possibility that the RI test itself might affect ACC structure, we also included animals (n = 6 per strain) in our analyses that were not subjected to the RI test. Neither measures of neuron death nor neuron density showed differences depending on whether animals were to the RI test, suggesting that the observed effects are not induced by our behavioural testing (Supp. Mat. Fig. 2a-b).

**Figure 1.**
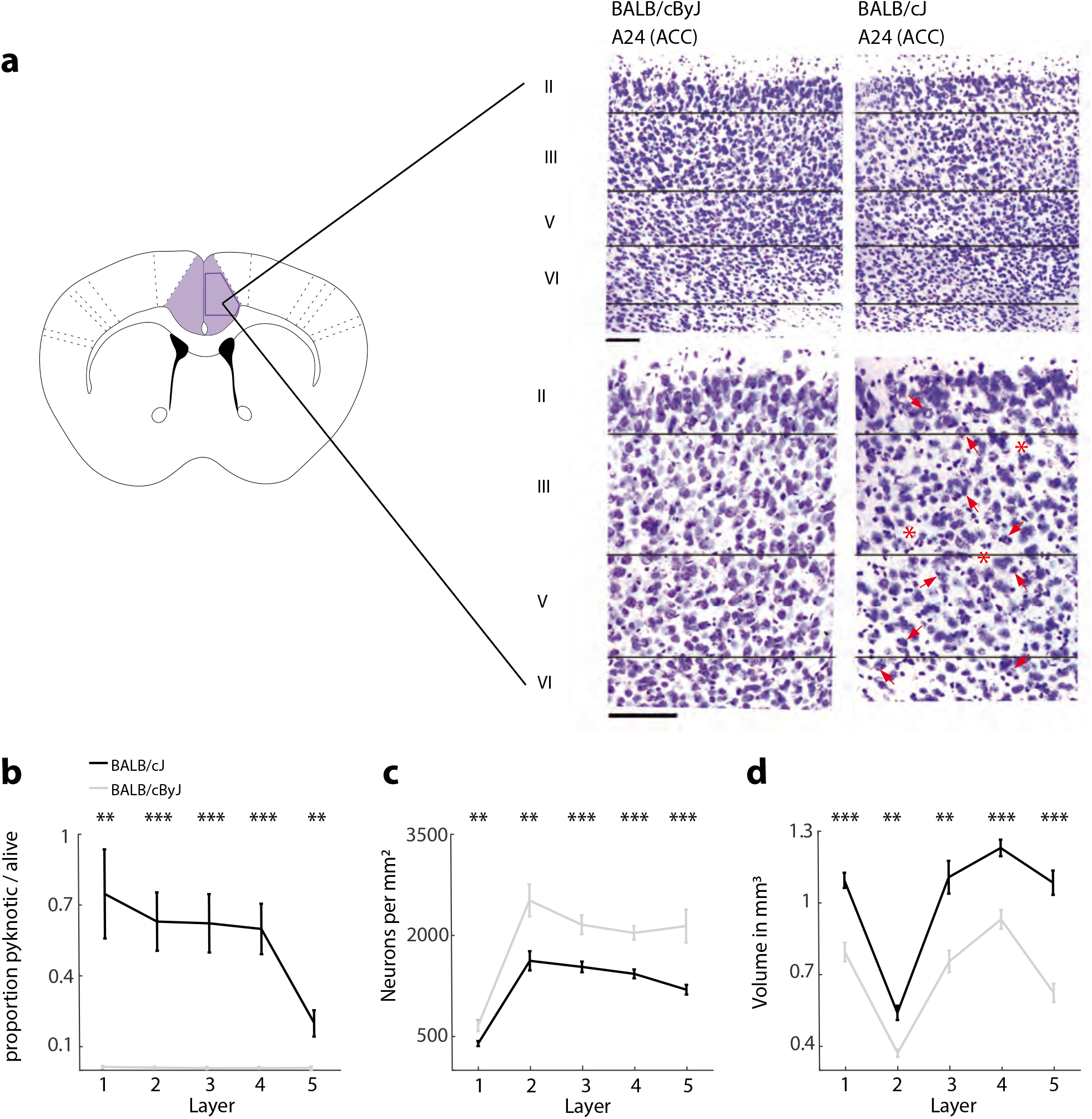
Cytoarchitecture of ACC**. a** Schematic of area 24 of ACC. Inset: Photographs in different magnifications showing the outlines of the position of layers. Pyknotic neurons are noted with red arrows in the high magnification BALB/cJ case. There are also 3 red asterisks to emphasize places where there are clearings with no neurons to emphasize their loss. Scale bars: 100 μm**. b** The proportion of pyknotic/non-pyknotic neurons. Black line: BALB/cJ mice, grey line: BALB/cByJ mice. Shown are average and SEM per layer**. c** The number of 23 neurons per mm^2^. Shown are average and SEM per layer**. d** The total volume in mm^3^. Black line: BALB/cJ mice, grey line: BALB/cByJ mice. Shown are average and SEM per layer. * p < .05, ** p < .01, *** p < .001.

To test if these measurements reflected a brain-wide phenomenon, we also measured pyknosis and neuron density in a neighbouring area: Secondary motor cortex (M2), which directly borders on ACC. While we also observed pyknosis here, there was no difference in neuron density (Supp. Mat. Fig 3a-b). This suggests that neuron degeneration only recently started in M2 and therefore has not yet affected neuron density. To further verify this trend, we performed a second staining directly targeted towards degenerating neurons (fluorojade C, [22]) and included somatosensory cortex (S1 mouth area, about 2.5 mm lateral distance to ACC) as well as insula (about 4.5 mm lateral distance to ACC) in the analysis. Again, we observed high levels of degenerating neurons in both ACC and M2 of BALB/cJ mice, but few in S1 and even less in insula. Together these results suggest that in BALB/cJ mice, neuronal degeneration begins in ACC, and then progressively spreads into neighbouring areas (Supp. Mat. Fig. 3c).

Interestingly, despite the dramatic amount of neuronal death and decreased neuronal density, across all layers of ACC, BALB/cJ mice showed an increased volume compared to BALB/cByJ mice (Fig. 1d, repeated measures ANOVA: *F*(1, 8) = 45.46, *p* < .001, *η2* = .85). This suggests that other cell types might be involved in extending or ‘bloating’ ACC.

### Increased neurotoxicity of astroglia aligns with neuronal death

Since both microglia and astroglia are activated by neuronal insult, we tested whether the widespread cell death in ACC had initiated abnormal glial processes and whether glial populations were generally increased. Therefore, we stained for microglia and astroglia, and then differentiated between dormant and reactive astroglia by determining the percentage of reactive toxic (A1) astroglia. In ACC, we observed a clear increase in the number of microglia (marker: Iba1) in BALB/cJ mice compared to control mice (Fig. 2a-b, repeated measures ANOVA: *F*(1, 27) = 7.79, *p* = .02, *η2* = .22). Post-hoc tests revealed that these differences were layer-specific (Fig. 2b): increases in microglia were restricted to layers 1 (post-hoc t-tests, p < .001) and 5 (p < .05). We next stained for the total number of astroglia (marker S100B), irrespective of their activity state, and observed increases in the number of astroglia in ACC (*F*(1, 26) = 9.12, *p* = .008, *η2* = .26) which were again layer-specific (post-hoc t-tests, layer 1 and 6, Fig. 2c-d). In a next step, we identified reactive astrogliosis by a GFAP staining, a marker that is often reported to be upregulated in reactive astroglia [23, 24] and observed dramatic increases in the number of GFAP-positive astroglia across all layers in BALB/cJ mice (Fig. 2e-f, repeated measures ANOVA: ACC: (*F*(1, 27) = 136.83, *p* < .001, *η2* = .84). In some layers, up to a ten-fold difference was observed, suggesting reactive astrogliosis may be responsible for the increased volume of ACC observed here and previously [18]. To decipher whether these reactive astroglia were toxic A1 astroglia, we performed a co-staining Serping1 (marker upregulated in A1 astroglia) with S100B (general marker for astroglia). Our data show high levels of neurotoxic A1 astroglia in BALB/cJ mice compared to BALB/cByJ mice in ACC (Fig. 2g-h, repeated measures ANOVA: *F*(1, 26) = 12.39, *p* = .003, *η2* = .32) and these effects were seen across all layers (post-hoc t-tests, all p < .001; Fig. 2D). This is in line with the high levels of pyknosis as A1 astroglia are known to induce cell death [25]. Again, as observed for neuronal death and density, there were no differences between mice that participated in the RI test and those that did not (Supp. Mat. Fig. 2c-f). Interestingly, we also did not observe differences in microglial and astroglial numbers in M2 and while we saw an increased number of reactive astroglia in M2, we did not see an increase in toxic A1 astroglia (Supp. Mat. Fig 3d-g). This again suggests that the structural degradation starts in cingulate cortex and might slowly further advance across cortex.

**Figure 2.**
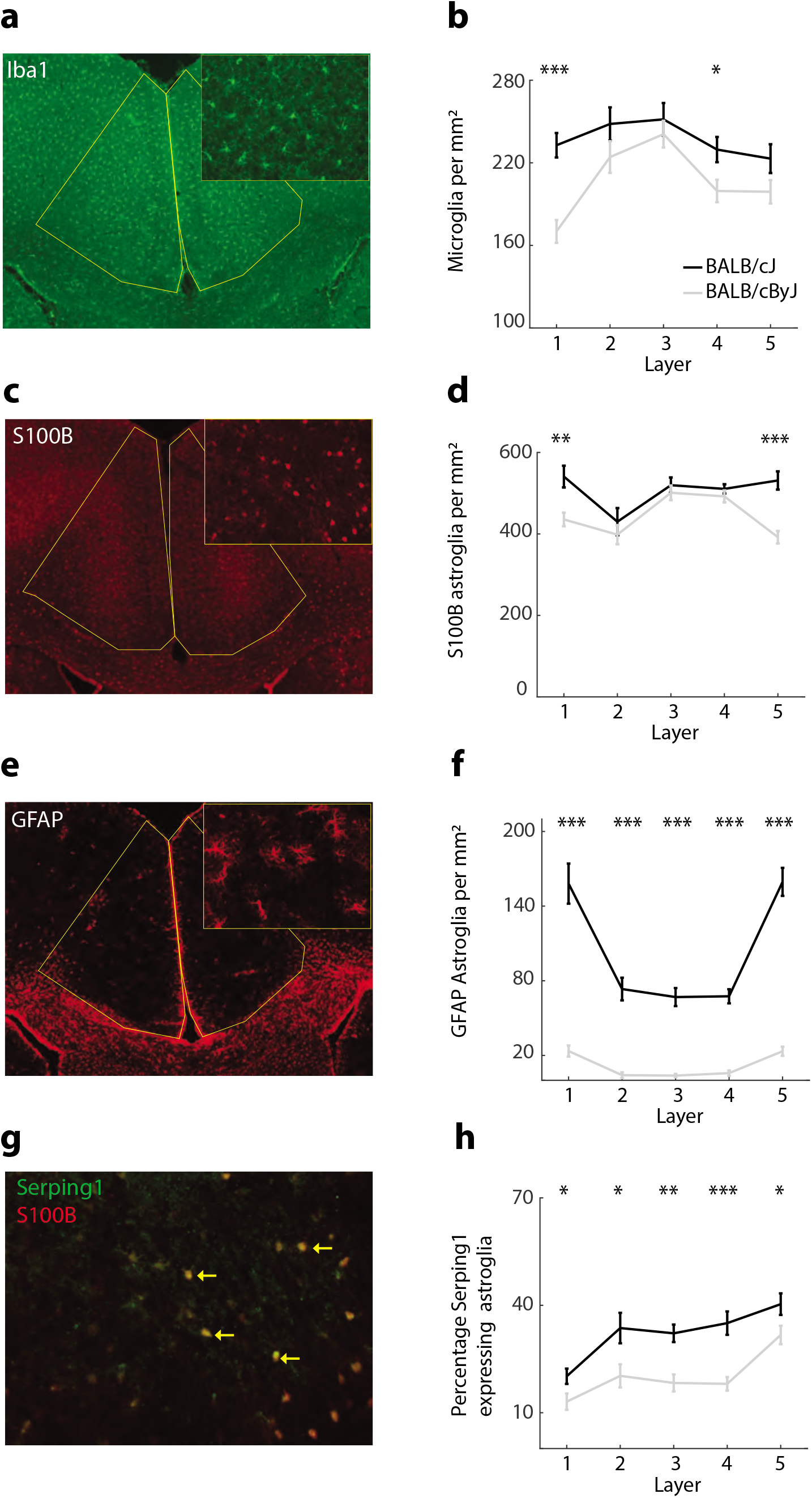
Microglia and astroglia across ACC layers. **a**. Example photograph of microglia across layers**. b** The number of microglia per mm^2^. Black line: BALB/cJ mice, grey line: BALB/cByJ mice. Shown are average and SEM per layer. **c-d** Same as a-b for S100B-positive stained astroglia across layers. **e-f** Same as a-b for GFAP-positive stained astroglia**. g** Example photograph of S100B astroglia double stained with a marker for toxic astroglia [Serping1, yellow arrow]. **h** The percentage of toxic astroglia. * p < .05, ** p < .01, *** p < .001.

### Gene-set enrichment analysis reveals genetic drivers of neuron death

We investigated whether gene-set enrichment analysis of genetic differences between BALB/cJ and BALB/cByJ strains, obtained from whole-genome sequencing data, would support our histological findings of neuronal death in the ACC. Single nucleotide variants (SNVs) that differed between the two strains were annotated to a total of 1541 genes, that were subsequently subdivided into 3 different categories: intronic/exonic non-coding and synonymous variants (1422 genes), untranslated region (UTR) variants (3’ & 5’, 90 genes), and missense mutations and splicing variants (61 genes). To analyse these genetic differences, we utilized the Ingenuity Pathway Analysis software package (IPA, QIAGEN). Four different analyses were performed: 1) IPA on genes annotated to the whole set of SNVs, 2) IPA on genes annotated to intronic/exonic non-coding/synonymous variants, 3) IPA on genes annotated to UTR variants, and 4) IPA on genes annotated to missense mutations and splicing variants. Data analysis was hypothesis-driven, and we report the results for the ‘cell death & survival’ category and its subcategories as provided by IPA (only categories that show at least 2 affected molecules).

#### All genes annotated to SNV differences in BALB/c strains (1514 genes)

IPA analysis of all SNVs supported our histological data: we found clear indications for genetic differences between BALB/cJ and BALB/cByJ mice related to cell death. Not only did IPA reveal differences in SNVs associated with apoptosis and necrosis but it specifically pointed towards degeneration of neurons and loss of brain cells (Table 1). As such, this is not only in line with the observed neuronal death but also the reduced neuronal density in ACC of aggressive BALB/cJ mice. Interestingly, we did not observe significant enrichment of categories related to glial involvement (after expanding the analysis to the nervous system category which contains the glial-related genes), suggesting that neuronal death is not a consequence of rogue glia but rather that glial processes are only activated upon neuronal death.

#### Genes annotated to intronic/exonic non-coding and synonymous SNV differences in BALB/c strains (1422 genes)

We subsequently looked at the intronic/exonic non-coding and synonymous variants separately, and again, the categories within cell death and survival specifically pointed to neuron degeneration and the loss of brain cells (Table 1).

#### Genes annotated to UTR SNV differences in BALB/c strains (90 genes)

IPA analysis of the UTR variants revealed SNV differences between the strains to be enriched for apoptosis of pericytes, erythroid precursor cells and erythroblasts. Even though degeneration of neurons was not highlighted in this analysis, the affected SNVs are also annotated to genes that overlap with genes from the whole dataset found to be involved in the degeneration of neurons.

#### Genes annotated to missense mutations and splicing region SNV differences in BALB/c strains (61 genes)

Similarly to the UTR analysis, the missense mutations and splicing variants did not show enrichment for neuron degeneration. The only significantly enriched category here was cytolysis of lymphocytes. Nevertheless, as with the UTR variants, the affected SNVs were also annotated to genes that were part of the category ‘degeneration of neurons’. The relatively low number of genes annotated to missense/splicing variants as well as UTR SNVs may, however, prohibit enrichment of the neuron degeneration category.

### Functional consequences of structural degradation

We hypothesised that the observed structural changes in ACC would have an impact on functional activation of ACC, and we therefore stained for an activity marker, cFos (Fig. 3a). At baseline, when animals had never been subjected to the RI test, there were no significant differences in ACC activity between BALB/cJ and BALB/cByJ mice (Fig. 3b, *F*(1, 10) = 1.26, *p* = .39, *η2* = .11). To track activity levels during aggressive encounters, we sacrificed animals 45-60 minutes after the last RI to allow for optimal cFos expression levels. Interestingly, after aggressive encounters, cfos markers revealed a lack of engagement of ACC in BALB/cJ mice: aggressive BALB/cJ mice activated markedly less cells compared to BALB/cByJ mice (Fig. 3c, left panel; *F*(1, 15) = 32.63, *p* < .001, *η2* = .69). This difference also held when we controlled for general neuron density across all layers of ACC (Fig. 3d, right panel; *F*(1, 15) = 28.34, *p* < .001, *η2* = .66). On average, BALB/cByJ mice activated more than twice the number of neurons during the RI test than BALB/cJ mice (Fig. 3c). As a consequence, ACC activity in BALB/cJ mice was not elevated after the RI test compared to baseline measurements (Fig. 3e, *F*(1, 13) = 0.95, *p* = .351, *η2* = .07), but clearly elevated in BALB/cByJ mice (Fig. 3f, *F*(1, 11) = 16.78, *p* = < .001, *η2* = .58). This suggests that while the structural degradation of ACC in BALB/cJ mice does not significantly impact resting ACC activity, it results in a failure to activate ACC during aggressive encounters. This in turn likely results in decreased top-down control of subcortical structures such as the amygdala, hypothalamus and PAG [26–29].

**Figure 3.**
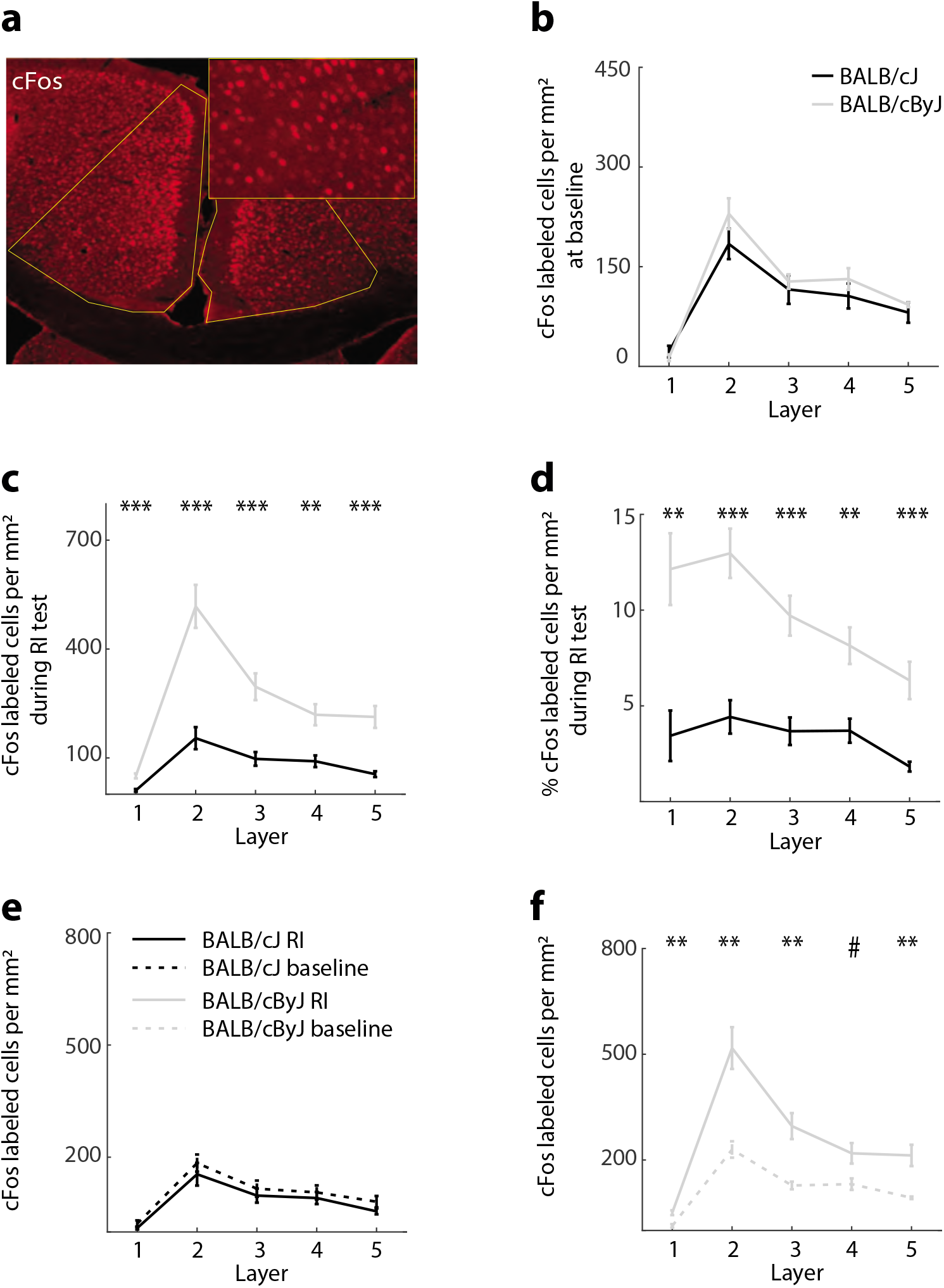
Activity levels. **a** Example photograph of stained cFos cells. **b** The number of cFos labelled cells in per mm^2^ at baseline. Black line: BALB/cJ mice, grey line: BALB/cByJ mice. Shown are average and SEM per layer. **c** The number of cFos labelled cells per mm^2^ during the last day of the RI test. **d** Same as b but expressed as percentage of total neuron density. **e** Comparison of activity at baseline vs RI for BALB/cJ mice. See in-figure legend. **f** Same as e for BALB/cByJ mice. * p < .05, ** p < .01, *** p < .001.

### Restoring ACC activity prevents escalatory aggression

Based on the observed hypofunction of ACC in BALB/cJ mice, we examined whether increasing ACC activity would be sufficient to prevent the escalation of anti-social aggressive behaviour. We chose to boost ACC activity chemogenetically: mice were injected bilaterally with an excitatory *designer receptor exclusively activated by designer drug* (DREADD, CaMKIIa promotor, hM3D(Gq receptor)) into A24 of ACC, while the control group only received a virus containing a fluorescent protein (control group). RI tests were performed two weeks after surgeries and all mice received clozapine-N-oxide (CNO) injections 30 minutes before testing. This was done to ensure that there are no off-target effects of the CNO.

Histology confirmed that virus expression was strong in A24 of ACC, with minimal bleeding into neighbouring M2 (Fig. 4a). Given that our previous experiments demonstrated that M2 is not specifically recruited during aggressive encounters, this minimal expression within M2 should not have had any behavioural effects. cFos histology confirmed that we indeed increased ACC activity in the DREADD group mice compared to the control group (Fig. 4b, *F*(1, 10) = 12.34, *p* = .01, *η2* = .53). On the behavioural level, we observed a marked decrease of aggression in the DREADD group (Fig. 4C): Mice attacked later (*F*(1, 10) = 11.77, *p* = .008, *η2* = .54) and showed fewer total bites (*F*(1, 10) = 39.8, *p* < .001, *η2* = .8; Fig. 5a). The bite pattern analysis (Fig. 5b) revealed that mice in the DREADD group performed fewer back bites (*F*(1, 10) = 20.6, *p* = .002, *η2* = .67) and fewer anti-social bites (*F*(1, 10) = 36.3, *p* = .009, *η2* = .78). Interestingly, the effect on the anti-social attack rate was much more pronounced than on typical back bites, and there was no difference in non-violent threat behaviour i.e. tail rattles (Fig. 5b, *F*(1, 10) = 2.7, *p* = .13, *η2* = .21). This suggests that increased ACC activity specifically curbs context-inappropriate anti-social aggression (Fig. 4D). Behavioural differences across individual sessions of the RI test can be found in Supplementary Figure 3. Correlation analyses demonstrated strong negative relationships between cFos markers of ACC activity during the RI-Test and all metrics of aggression except for threat behaviour (tail rattles): The higher ACC activity, the less physical aggression was observed (Fig. 5c-d; all R^2^ > .45, all p < .02 after correction for multiple comparisons).

**Figure 4.**
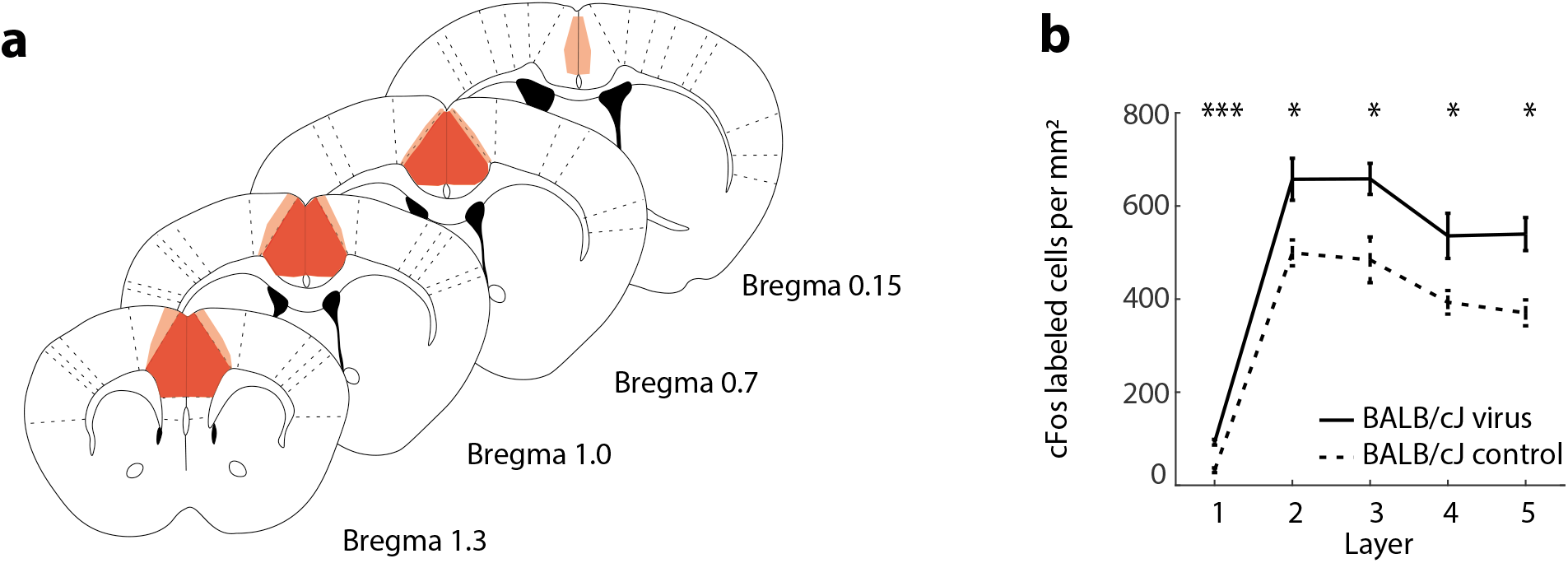
Chemogenetic manipulation of ACC activity. **a** Verification of virus expression, bright red = strong expression, light red = sparse expression. **b** Average of cFos-labelled cells across ACC layers. Solid line: DREADD group. Dashed: Control BALB/cJ mice. Error bars: SEM.

**Figure 5.**
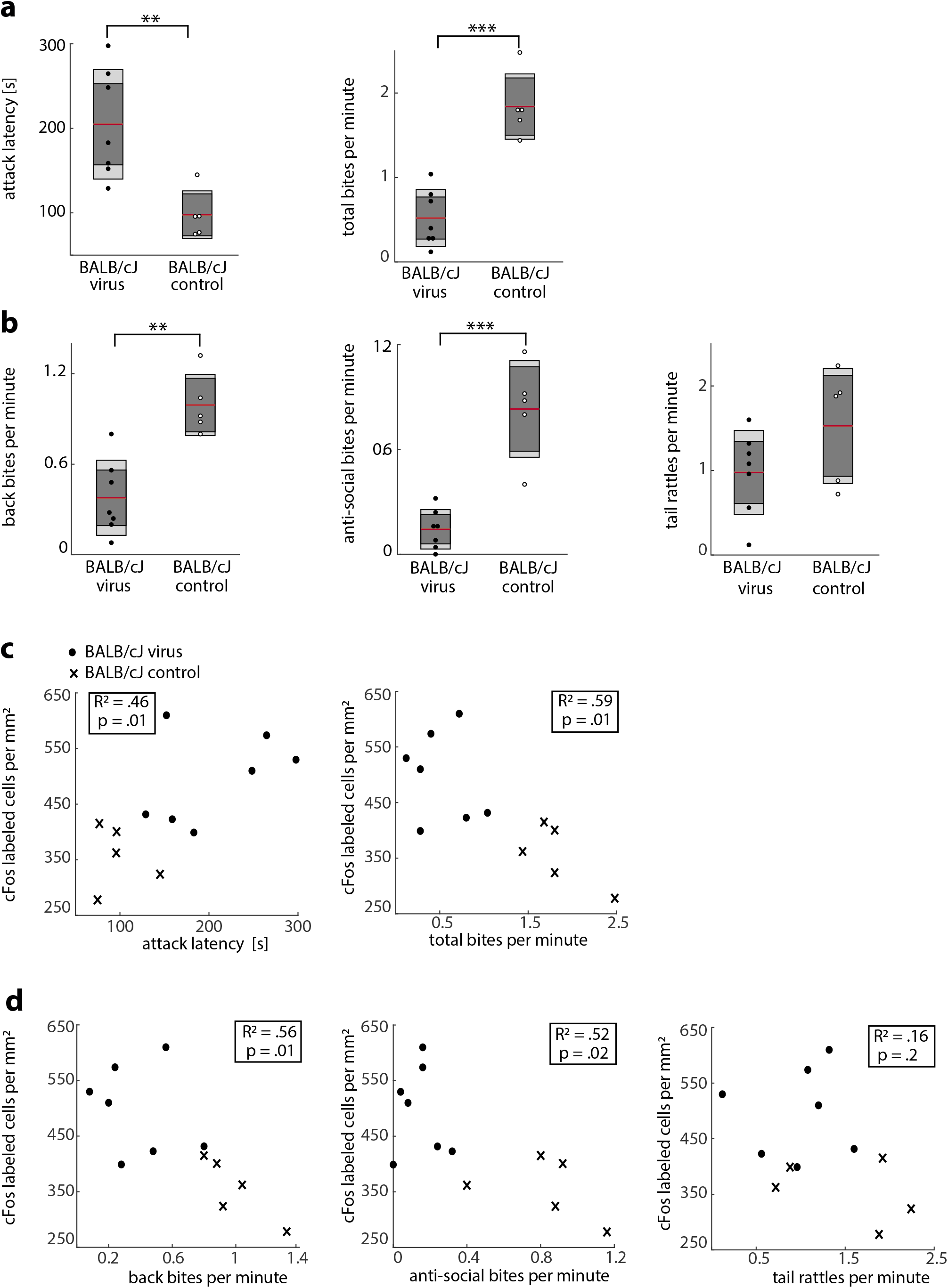
Effect of chemogenetic manipulation on aggressive behaviour. **a** Average attack latency (left), average total bites per minute (right). Black dots: BALB/cJ mice injected with chemogenetics virus group, white dots: BALB/cJ mice injected with control virus group. Red line: Average; Dark grey area: 67% confidence interval; Light grey area: 95% confidence interval. **b** Same as a for back bites (left), anti-social bites (right) and tail rattles (right). **c** Correlations between average ACC activity and attack latency (left) and total bites (right). Dots: BALB/cJ mice injected with DREADD virus; Crosses: Control BALB/cJ mice. **d** Same as c for back bites (left), anti-social bites (middle) and tail rattles (right). * p < .05, ** p < .01, *** p < .001.

To understand how ACC activity was able to specifically curb anti-social aggression, we probed its effect on downstream structures by applying cfos staining to prominent efferent regions, specifically the basolateral amygdala (BLA), the lateral hypothalamus (LH), the ventromedial hypothalamus (VMH) and the mediodorsal thalamus (MD). BLA, LH and VMH are considered part of the threat circuit [30, 31] and MD has been associated with flexible goal-directed behaviour [32, 33]. We saw direct efferent projections from ACC to BLA, LH and MD but none to VMH (Fig. 6a). However, since LH is known to project to VMH [34], we decided to include VHM in our analyses as a disynaptic target region. For BLA, LH and VMH we observed substantial decreases in activity in the DREADD group compared to the sham virus group (all p < .05, Fig. 6b; see Supp. Mat. Fig. 5 for activity per sub-area of these structures). Interestingly, when correlating activity measures within these regions to the behavioural metrics, we observed that activity within BLA, LH and VMH positively predicted the amount of bites (Fig. 6c; all R_2_ > .35; all p < .05 after correction for multiple comparisons), but only BLA also predicted attack latency (R_2_ = .50, p = .04; all other R2 < .35, all other p > .1; see Supp. Mat. Fig. 6). This suggests that these downstream regions jointly determine the level of aggressive behaviour (i.e. the amount of bites), but within only BLA seems to encode the initial drive to attack. For MD, we observed a negative correlation with behaviour: Higher MD activity led to fewer bites (R^2^ = .56, p = .02), but did again not affect attack latency (R^2^ = .32, p = .18).

**Figure 6.**
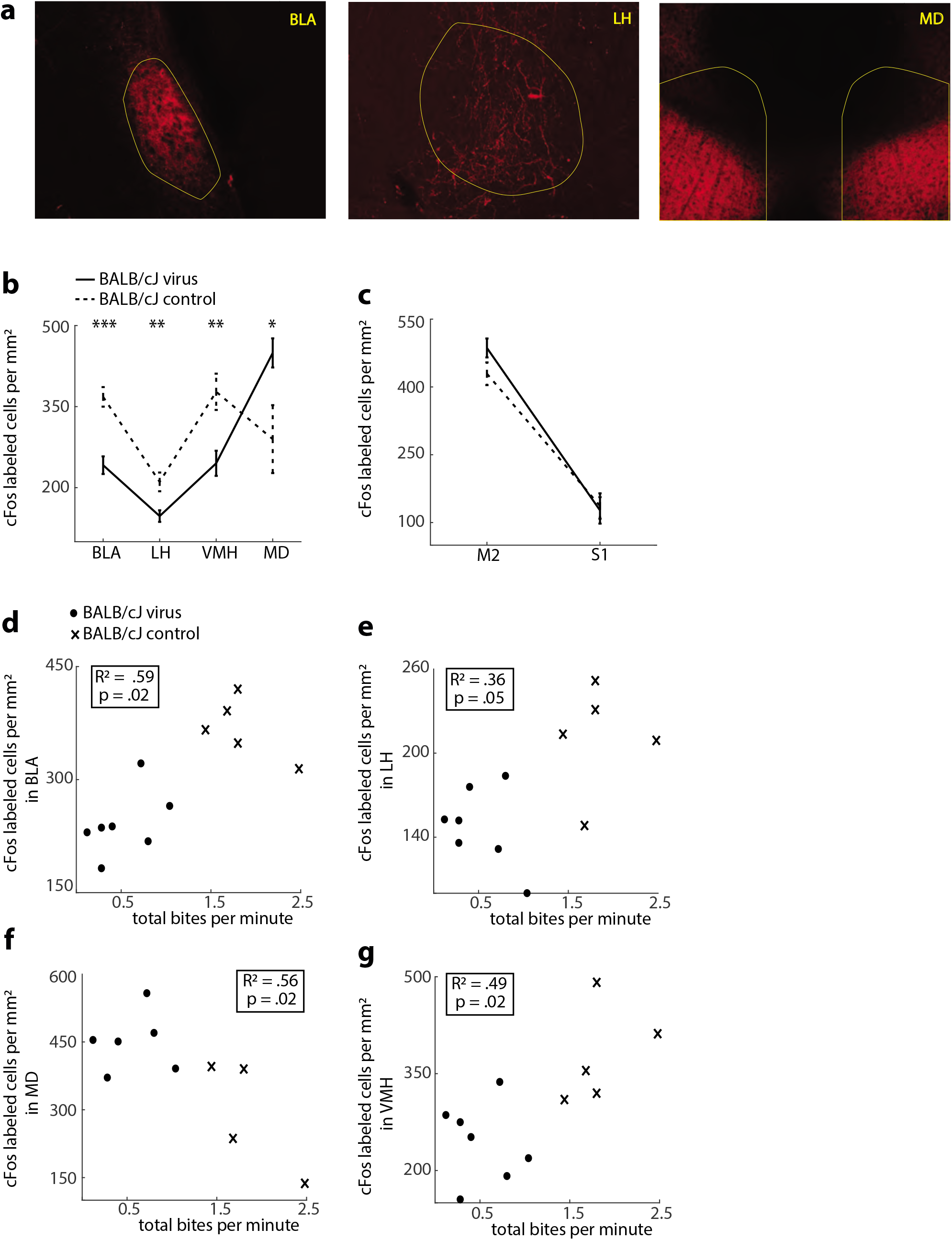
cFos activity in downstream regions of ACC. **a** Left panel: Efferents of ACC in basolateral amygdala (BLA), lateral hypothalamus (LH) and mediodorsal thalamus (MD). **b** Activity within subcortical regions. Black line: BALB/cJ mice injected with chemogenetic virus, dashed line: BALB/cJ mice injected with control virus. Shown are average and SEM. **c** Same as b for activity within M2 and S1. **d** Correlation between BLA activity and average total bites per minute. Black dots: Mice injected with chemogenetic virus, crosses: mice injected with control virus. See in-figure legend. **e** Same as d for LH. **f** Same as d for MD. **g** Same as d for VMH. * p < .05, ** p < .01, *** p < .001.

To control whether our chemogenetic manipulation may have resulted in a non-specific activation of cortical activity, we also checked cFos expression in M2 and S1 in the DREADD group. We did not observe any differences in activity here, neither for M2 (Fig. 6c, ANOVA: *F*(1, 10) = 3.07, *p* = .13, *η2* = .26) nor S1 (ANOVA: *F*(1, 10) = 0.5, *p* = .82, *η2* = .005). This supports the notion that our manipulation was indeed confined to ACC, which then resulted in strong and specific downstream network effects.

## Discussion

Here, we demonstrate a causal role for ACC in preventing anti-social aggression in BALB/cJ mice. We show that the ACC of pathologically aggressive BALB/cJ mice is structurally degraded, exhibiting extensive neuron death coupled with a drastic loss of total neuron density. In line with this, glial populations showed an increase in the number of microglia and astroglia as well as neuro-toxic astrogliosis. Genes annotated to SNV differences between BALB/cJ and BALB/cByJ strains showed enrichment for neuronal degeneration, further confirming the notion that neuron death drives astrogliosis. Importantly, cell death was not a brain-wide phenomenon, but seemed to focus on ACC, and gradually spread outwards to neighbouring areas such as M2. The structural degradation of ACC had functional repercussions: Unlike control mice, BALB/cJ mice failed to activate ACC during aggressive encounters. By heightening ACC activity chemogenetically, we were able to prevent escalatory aggression, particularly anti-social aggression. Analysis of cfos expression in downstream regions suggested that this effect was likely due to downregulation of ‘threat circuit’ regions such as BLA, LH and VMH, and upregulation of activity in MD.

Our observation of neuron death raises several questions. First, does the involvement of glial populations and toxic astrogliosis imply that neuronal insult initiates a cascade of glial responses or that glial processes initiate neuron death? Our data indicate that neuronal insult comes first. The gene-set enrichment analysis based on SNV differences between the two BALB strains did not point to glia, but highlighted several genes connected to neuronal degeneration. In addition, in neighbouring M2 we observed neuron death but not toxic A1 astroglia (Supp. Mat. Fig. 2). This suggests that neuron death starts in ACC due to neuronal insult (e.g. based on developmental factors) and this recruits glial processes to remove neuronal debris. At this point, toxic astrogliosis likely starts, which in turn accelerates neuron death.

Another question is whether cell death affects specific neuronal populations differentially. Although we did not label individual neuron types in this study, there is some evidence that this is unlikely to be the case. First, we have previously demonstrated that the density of PV and SOM interneurons decreases across cingulate cortex in BALB/cJ mice [19]. Our current results would suggest that this decrease in interneuron density is the product of neuron degeneration across ACC. In the previous study [19], we did not measure the density of excitatory neurons across cingulate cortex. However, given that interneurons only represent about 10% of the neuronal population, the fact that we observed extensive neuron degeneration across all ACC layers (Fig. 1B) suggests that not just interneurons but also principal neurons are degenerating in approximately equal measure. This is also in line with our finding that upregulating the remaining population rescues normal aggression control (see below), which suggests that cell death has not erased specific populations or connections.

Based on the decreasing rates of neuron death observed across neighbouring brain regions, it appears that structural degradation begins in cingulate cortex and then gradually spreads to neighbouring areas. Why would cingulate cortex be the ‘epicentre’ of cell death? The key may lie in a shift of developmental processes: While neuron death occurs naturally across all brain areas during development, cingulate cortex is known to undergo a particularly high level of cell death [35, 36]. This means that if the stop signal for developmentally necessary neuron death was to fail, continual pruning would have particularly devastating effects on cingulate cortex. The spread of neuron death to neighbouring areas (Supp. Mat. Fig. 2A, right panel) could then be explained by a cascade of neuronal and glial signaling: A1 astroglia lose their ability to promote neuronal survival and synaptogenesis and instead transmit a potent neurotoxin that rapidly kills neurons [25]. Such dying neurons can then induce a process called transneuronal degeneration or secondary neuron loss: neurons connected to those dying might die themselves, either due to the loss of synaptic input (anterograde transneuronal degeneration) or due to the loss of synaptic outputs (retrograde transneuronal degeneration) [37]. Our mice were tested as young adults (10-11 weeks old). If the hypothesis stated here is true, older BALB/cJ mice should show increasing levels of neuronal degeneration across cortex.

An additional mechanism that might exacerbate neuron death in cingulate cortex is a change in the GABAergic switch during development. During early development, GABAergic transmission is excitatory and only switches to being inhibitory during the early postnatal period [38]. Recent work demonstrated that BALB/cJ mice have increased expression of the Kcc2 gene [39] which encodes a protein that is crucial for correct GABAergic switching. If this switch is delayed, excessive excitation might induce neuron death via excitoxicity and oxytosis [37].

One of the particularly interesting features of our chemogenetic perturbation was that simply increasing ACC activity was enough to induce a behavioural rescue – the BALB/cJ mice that received the DREADD virus showed virtually identical behaviour to control mice. Our results suggest that this does not require a specific pattern of ACC activity, but rather a simple augmentation of ACC output. This is surprising because the behavioural change induced by this manipulation is quite intricate: Anti-social aggression is strongly suppressed while context-appropriate aggression is reduced less strongly (see next paragraph). While surprising, this type of effect is consistent with studies in other brain regions, particularly hippocampus, which show that complex cognitive processes and behavioural responses (e.g. the recall of contextual memories) can be triggered by simple activation of the neuronal populations involved [40].

As mentioned above, the activation of ACC had a clearly differential effect on aggressive behaviour: Anti-social biting, which is normally a defining behavioural feature of BALB/cJ mice, was almost entirely eradicated – just as in control mice. In contrast, context-appropriate bites were reduced less dramatically and threat behaviour remained constant. Interestingly, during the last two days of the RI test (Supp. Mat. Fig. 3), the mice in the DREADD group started to show increased socially expected biting and threat behaviour – again very similar to control mice. These increases can be interpreted as an adaptive aggressive response to repeated territorial challenge [18, 19]. This suggests that boosting ACC activity removed the anti-social aspects of aggression, but left adaptive aggression largely intact. Based on their threat behaviour and context-appropriate biting, it stands to reason that animals in the DREADD group still experienced the RI test as threatening; however, they were now able to respond context-appropriately by refraining from anti-social attacks.

These results further clarify the role of ACC in the control of aggressive behaviour. It was previously shown that inhibiting ACC in wildtype mice during aggressive encounters leads to sudden bursts of aggressive behaviour [14]. This can in principle indicate that ACC activity curtails aggressive behaviour in general. Here we show that rather than producing a blanket inhibition of aggressive behaviour, ACC activity seems to selectively ‘edit’ aggressive impulses, inhibiting mainly context-inappropriate expressions of aggression. When faced with an intruder, a warning consisting of tail rattles or a few harmless attacks to the back is context-appropriate, whereas potentially lethal bites to the abdomen or neck represent a maladaptive strategy. By activating ACC in BALB/cJ mice, it appears that we were able to re-instate ACC’s ability to assess the situation and adapt the aggressive reaction accordingly.

How is such ‘editing’ of aggressive impulses by ACC accomplished? Our findings suggest that ACC activity ‘fine-tunes’ the balance of activity in downstream regions: On the one hand, activating ACC increased activity in MD – a connection that has previously been shown to contribute to behavioural flexibility [31, 32]. On the other hand, restoring ACC activity led to a suppression of subcortical regions, particularly BLA, LH and VMH. All three of these regions form important parts of the threat circuit (Fig. 5). BLA appears to encode the initial drive to attack, as activity within BLA correlated negatively with attack latency, while LH and VMH activity correlated only with the number of bites. This also explains to some extent how antisocial aggression can be reduced selectively by targeting downstream regions differentially: For instance, ACC projections to LH have previously been shown to be specifically involved in anti-social aggression [26] whereas projections to mediobasal hypothalamus seem to affect the general incidence of aggressive behaviour.

Our results have several implications for understanding the mechanisms of maladaptive aggression. First, previous work has shown that even when one increases the drive to attack by manipulating activity within VMH [8], context-sensitive control mechanisms are still in place, as certain contextual cues (e.g. presence of a female mouse) reduced aggression even under optogenetic stimulation. Our work seems to explain why: ACC utilises these contextual cues to edit aggressive drives. When ACC activity fails, as we observe in our BALB/cJ mice, the drive to attack cannot be modulated and results in escalatory aggression and anti-social attacks.

Reduced activity of ACC has also been observed in human patients with pathological aggression [2, 13] but there is little consensus on the underlying structural alterations. Several studies demonstrated reduced volumes of ACC while others reported increased ACC volume [15, 41]. Our data might resolve this discrepancy: as neuron degeneration starts, glial populations will start to massively proliferate which might explain increases in volume. However, as soon as degenerated neurons are ‘cleaned up’, the volume might be reduced due to the neuron loss and the reduction of glial activity. Post-mortem studies of ACC of patients with pathological aggression would be needed to investigate whether neuronal degeneration and toxic astrogliosis can be observed in human patients. Our functional data also suggest ACC as a prime target for clinical interventions in dysfunctional aggression: Based on our results, it appears that increasing ACC activity, for example via transcranial magnetic stimulation, may be able to address maladaptive aggression with minimal side effects, leaving context-appropriate aggression and patients’ overall affect intact.

In conclusion, to our knowledge, we are the first to provide a causal link between ACC activity and the control of escalatory and anti-social aggression. We show that ACC activity shapes dynamics in subcortical circuits and that re-instating ACC activity seems to establish sufficient top-down control over these circuits to all but eliminate anti-social aggression. At the same time, our data provide important insights into the pathology of aggression and suggest structural degradation as underlying mechanism of altered ACC activity in aggression.

## Material & methods

### Behaviour

#### Animals & Housing conditions

For all experiments, BALB/cJ and BALB/cByJ mice were obtained obtained from Jackson Laboratory (Bar Harbor, ME, USA) and C57BL/6J intruder mice were obtained from Charles River Laboratories (Erkrath, Germany). All mice were housed in an enriched environment (High Makrolon^®^ cages with Enviro Dri^®^ bedding material and Mouse Igloo^®^) and had free access to dry food and water. They were kept at a reversed 12–12 h day–night cycle with sunrise at 7.30 pm. In line with the typical resident intruder (RI) protocol, test mice were housed individually, while intruder mice were housed in groups of 5–6 animals per cage. At the start of the RI test, all resident BALB/cJ and BALB/cByJ mice were 11 weeks old and all intruder C57BL/6J mice were 7 weeks old. All animal procedures were conducted in compliance with EU and national regulations as well as local animal use ethical committees (European Directive 2010/63/ EU), and approved by the Ethics Committee on Animal Experimentation of Radboud University (RU-DEC number 2013-235 & RU-DEC number 2017-0032).

#### RI test

Resident mice were housed individually 10 days prior to testing. In total we tested 15 BALB/cJ mice and 13 BALB/cByJ mice across 3 different cohorts to account for possible litter effects. Aggression testing was done in the home cage of the BALB/cJ and BALB/cByJ mice in a dark room with red overhead lighting. Behaviour was videotaped using an infrared camera (SuperLoLux, JVC). Animals were tested for five consecutive days, and each day each BALB/cJ and BALB/cByJ mouse was confronted in their home cage with a different C57BL/6J intruder mouse. The order of testing was randomised and the first test was started 1 hour after the beginning of the dark phase (active phase). Testing started by placing an intruder animal in the home cage of the resident animal, separated by a glass screen to allow for visual and olfactory stimulation for 5 min. After this instigation phase, the glass screen was removed and interaction was allowed for 5 minutes. In addition, we subjected 5 BALB/cJ and 5 BALB/cByJ mice (from here on referred to as cohort 3) to a slightly different protocol: we allowed interaction for 5 min *after* the first attack (up to a total maximum of 10 min). This was done to exclude the confound that BALB/cByJ mice might need longer time to show aggression as a late attack latency might influence the comparison of the total number of bites between BALB/cJ and BALB/cByJ mice. By allowing 5 min of confrontation after their first attack, all mice in this group therefore had the same amount of time to show aggressive behaviour. Our results show that a prolonged confrontation phase made no difference for the BALB/cByJ mice, they were still significantly less aggressive than BALB/cJ mice also when compared to those that were tested in the shorter protocol (from here on referred to as cohort 1 & 2).

#### Data analysis RI test

All behavioural measures (except attack latency) are expressed as observation per minute such that the behaviour can be directly compared across cohorts. Attack behaviour was scored manually in terms of attack latency, attack frequency and tail rattles using the program The Observer (Noldus). An attack was defined as a bite directed at the back, tail, abdomen, flank, neck or face of an intruder [18–20]. Attacks directed to the abdomen, flank, neck or face are considered as anti-social bites as they have the potential to really inflict harm on the intruder [18, 19]. All recordings were scored by the same researcher who was blind to the strain of the animal (BALB/cJ and BALB/cByJ mice have the same appearance).

#### Statistical analysis

Aggressive behaviour was analysed with a repeated measures ANOVA (days as within-subject factor, strain as between-subject factor) to illustrate the course of aggressive behaviour across the 5 days of RI testing. Next, aggression scores were pooled across all days and then analysed with a one-way ANOVA (strain as between-subject factor). Attack latency was analysed separately for cohort 1-2 and cohort 3 to demonstrate that a longer confrontation time did not result in increased aggression in control mice. All other behavioural measures were analysed together for all cohorts. All statistical analyses were carried out using SPSS23-software (SPSS inc., Chicago, USA). The false discovery rate method [42] was used to correct for multiple comparisons for all behavioural data.

#### Perfusion and tissue preparation

After the last RI test, all BALB/cJ and BALB/cByJ mice were deeply anesthetized with isoflurane (3–5%) and perfused with saline, followed by 150 mL of 4% paraformaldehyde solution (PFA) in 0.1M phosphate buffer (PBS). Note: mice of cohorts 1 and 2 were perfused 45 to 55 minutes after the last RI confrontation to allow for immediate early gene expression analyses. Brains were removed, fixed overnight in 4% PFA and then kept in 0.1M PBS at 4-degree temperature. 1 day before cutting, brains were placed in 0.1 M PBS plus 30% sucrose to ensure cryoprotection. Coronal sections (30 μm) were obtained on a freezing microtome (Microm, Thermo Scientific). All sections containing ACC and MCC were placed in running order in containers filled with 0.1M PBS + 0.01% sodium azide (to prevent fungal contamination) and stored at 4-degree temperature until use.

### Cytoarchitecture ACC

#### Neuronal density & cell death across layers

To determine neuronal density and cell death we analysed ACC sections of 20 BALB/cJ mice and 13 control BALB/cByJ mice. Note: of the 20 BALB/cJ mice analysed, 6 mice did not perform the resident intruder test to control for any possible effects on density or cell death and ensure these were not the consequence of the RI test. As performed for the volume layer measurements, we used sections stained with cresyl violet and constructed contours of the individual layers of ACC. From each mouse the ACC sections were chosen at approximately the same anatomical level (ACC sections at AP 0.85). To localize A24, we used the Paxinos & Franklin mouse atlas [43] and relied on the same methodology as [18] to define the start/end of each area and the border with A24. The counting was done using the optical fractionator tool in the program StereoInvestigator (MBF Biosciences), as this represents an unbiased method for cell counting. Each section had the same counting frame (50×50) and grid size (100×100). Counting was done at a 60x magnification. The *estimated population using number weighted section thickness* value given by the program was used as value for neuronal density. First, the area of each layer was delineated and then the neurons in each layer counted. Cells with a distinct nucleolus with or without several heterochromatin granules and a rim of cytoplasm around the nucleus were considered neurons. Those without a distinct nucleolus were not considered neurons. In addition, we counted those neurons showing signs of cell death. These were defined as cells with an irregular, shrunken and/or crenulated shape and nuclear shrinkage as well as increased vacuolation and tissue disruption. The number of neurons was then calculated as the sum of both healthy and pyknotic neurons per mm^2^ (dividing the value provided by the program for each individual layer by the surface area of that layer). To determine the proportion of healthy neurons vs pyknotic neurons, we divided the number of pyknotic neurons by the number of healthy neurons. Data was analysed with repeated measures ANOVAs (layer as within-subject factor, strain as between subject factor) and t-tests were used as post-hoc tests. The false discovery rate method [42] was used to correct for multiple comparisons.

#### Volumetric layer measurements

For layer measurements, 5 BALB/cJ and 5 BALB/ cByJ mice previously tested in the RI test were used. As done in [18], we mounted all sections containing A24 (ACC) on gelatine-coated slides [0.5% gelatine + 0.05% potassium chromium (III) sulphate]. Sections were then air-dried and placed in a stove at 37° overnight. The next day, sections were first placed in a 96% alcohol bath for 10 min, then hydrated in graded alcohol baths (1 × 70%, 1 × 50%, 2 min each), dehydrated in graded alcohol baths (1 × 70%, 1 × 96%, 1 × 100%, 2 min each) and stained in a 0.1% cresyl violet solution for approximately 5 min.

Afterwards, sections were placed in a graded alcohol series (3 × 95%, 3 × 100%, 2 min each), cleared in xylene (Sigma–Aldrich) and mounted with entellan (Sigma–Aldrich). One image of each section was then obtained on an Axioskop fs microscope using Neurolucida software (MBF Bioscience). To localize A24 we used the Paxinos & Franklin mouse atlas [43] and relied on the same methodology as [18] to define the start/end of each area and the border with A24. Using Neurolucida, we then constructed contours for each cortical layer of each section separately. After constructing the contours for every ACC section, the volume per section was determined according to the following formula: area in mm_2_ × section thickness in mm = volume in mm^3^. Finally, A24 volume was computed by adding up the volumes of all relevant sections. All contours were drawn by the same researcher and the researcher was blind to the group of the animal to account for possible biases. The volume of the different layers of both A24 was then analysed with a repeated-measures ANOVA with strain as between-subject factor and layer as within-subject factor. T-tests for independent samples were then performed as post-hoc tests. All statistical analyses were carried out using SPSS23-software (SPSS Inc., Chicago, USA). The false discovery rate method [42] was used to correct for multiple comparisons.

### Glial changes in ACC

#### Microglia and astroglia across layers

To assess the number of microglia and astroglia, we used ACC sections of 15 BALB/cJ and 13 BALB/cByJ mice. As for the neuron density and pyknosis, for both strains we included animals (N = 6 per strain) that did not perform the RI test to exclude effects of the RI test on glial changes. ACC sections were stained with a standard free-floating immunofluorescence protocol with antibodies for microglial and astroglial markers (Iba1 and GFAP and S100B respectively). Briefly, sections were incubated overnight at room temperature in Iba1anti-rabbit (1:1500, Wako, product code: 019-19741) and GFAP anti-guinea pig (1:1500, Synaptic Systems, product code: 173004) as well as a marker for neurons (NeuN anti-chicken, 1:1000, Millipore, product code: ABN91). The following day, the sections were incubated in matching secondary antibodies at room temperature (Alexa Fluor donkey anti-rabbit 488 [Abcam, product code: ab150061], Alexa Fluor donkey anti-guinea pig 647 [Jackson Immuno Research, product code: 706-605-148] and Alexa Fluor goat anti-chicken 555 [Thermo Fisher, product code: A-21437]). Photographs of the ACC and MCC sections were then taken with an Axio Imager.A2 microscope (Zeiss) at 20x magnification and analysed with Fiji (ImageJ). In short, we used the NeuN stain to outline the different layers, saved these as ROIs and then applied them to the Iba1 and GFAP stains. Microglia and astroglia were then manually counted and their number divided by the surface area of the layer to attain the number of glia per mm^2^. We then analysed the data with repeated ANOVAs (layer as within-subject factor, strain as between subject factor) and t-tests were used as post-hoc tests. The false discovery rate method [42] was used to correct for multiple comparisons. Given that GFAP in grey matter is known to often preferentially stain reactive astroglia [24], we decided to stain for another astroglia marker, S100B, in combination with a marker for reactive toxic astroglia (Serping1). This enabled us to determine the total number of astroglia regardless their activity state as well as the percentage of toxic and neuroprotective reactive astroglia. As done previously, we used a standard free-floating immunofluorescence protocol and incubated the ACC and MCC sections overnight in S100B anti-guinea pig (1:1000, Synaptic Systems, product code: 287004) together with Serping1 anti-mouse (Santa Cruz, product code: sc-377062) as a marker for neurotoxic A1 astroglia. The next day, sections were incubated with matching secondary antibodies (anti-rabbit and anti-guinea pig were the same ones as previously used as well as an Alexa Fluor goat anti-mouse 555 [Abcam, product code: ab150114]) and a DAPI stain was added. Photographs of the ACC sections were then taken with an Axio Imager.A2 microscope (Zeiss) at 20x magnification and analysed with Fiji (ImageJ). Layers were outlined with the DAPI stain, ROIs saved and applied to the S100B, and Serping1 stains. Positively stained cells were then manually counted and Serping1 markers were overlayed with the S100B markers to count the number of double-stained cells. The number of S100B positive cells was divided by the surface area of the specific layer and the number of A1 astroglia was determined by calculating the percentage of S100B + Serping1 double-stained cells. We then analysed the data with repeated ANOVAs (layer as within-subject factor, strain as between subject factor) and t-tests were used as post-hoc tests. The false discovery rate method [42] was used to correct for multiple comparisons.

### Whole-genome sequencing and gene-set enrichment analysis based on genetic differences between the BALB strains

Sequencing libraries were prepared from high-quality genomic DNA using the TruSeq DNA PCR-Free kit (Illumina) and ultra-deep whole genome sequencing (average 30X read-depth across the genome) was performed on a HiSeq X Ten System (Illumina). We developed an efficient data processing and quality control pipeline. Briefly, raw sequencing data underwent stringent quality control and was aligned to either the mm10 [BALB/cJ vs BALB/cByJ strain comparison]. Isaac was used to align reads and call single nucleotide variations (SNVs). We excluded SNVs that were covered by less than 20 reads, and that were not present in both animals from the same strain. SnpEff was used to annotate SNVs and explore functional effects on gene function. As further described in the Results section, SNVs differing between the two strains were annotated to a total of 1541 genes, which were subdivided into 3 different categories (intronic/exonic non-coding and synonymous variants [1422 genes], UTR [3 & 5, 90 genes], missense mutations and splicing variants [61 genes]). These sets of genes were analysed with IPA: based on information from the published literature as well as gene expression and gene annotation databases, IPA assigns genes to different groups and categories of functionally related genes [44]. Given our histological data demonstrating neuronal death in ACC, we decided to test hypothesis-driven and focus on the category ‘cell death and survival’. IPA calculates p-values for the enrichment of each gene category using the right-tailed Fisher’s exact test. In addition, the Benjamini-Hochberg correction is used to account for multiple testing.

### Functional activation in ACC

To assess the number of active cells, we used ACC brain sections of 15 BALB/cJ and 13 BALB/cByJ mice. As previously, for both strains we included animals (N = 6 per strain) that did not perform the RI test to check for baseline activity of ACC. ACC sections were stained with a standard free-floating immunofluorescence protocol with an antibody for cFos. Briefly, sections were incubated overnight at room temperature in cFos anti-guinea pig (1:1000, Synaptic Systems, product code: 226004). The following day, the sections were incubated in a matching secondary antibody at room temperature (Alexa Fluor donkey anti-guinea pig 647) and a DAPI stain was added. Photographs of the ACC sections were then taken with an Axio Imager.A2 microscope (Zeiss) at 20x magnification and analysed with Fiji (ImageJ). Layers were outlined with the DAPI stain, ROIs saved and applied to the cFos stain. Positively stained cells were then manually counted. The number of cFos positive cells was divided by the surface area of the specific layer. We then analysed the data with repeated ANOVAs (layer as within-subject factor, strain as between subject factor) and t-tests were used as post-hoc tests. The false discovery rate method [42] was used to correct for multiple comparisons.

### Manipulating ACC activity

#### Surgery

Fifteen 8-week old BALB/cJ mice were used for this experiment. All mice underwent stereotactic surgery and received a viral injection. Briefly, mice were anaesthetized with a ketamine-dextomidor mix and bilaterally a small hole was made at the following coordinates: 1.0 AP, 0.3 ML and a viral construct was injected at a depth of 1.3 DV. 10 mice received a bilateral 1μl injection of a viral DREADD construct (pAAV-CaMKIIa-hM3D(Gq)-mCherry, Addgene, catalogue number: 50476-AAV5) to enable activation of infected neurons. The other 5 mice were injected with a fluorescent protein to mimic inflammation and surgical procedures and used as a control group. Mice were housed individually after surgery and testing was performed 3 weeks after surgery to allow for recovery and optimal viral expression.

#### Behaviour

Mice were tested in the RI test for 5 consecutive days (see Behaviour: Methods & Results, section RI test). 30 minutes before the start of the test each day, all mice received an i.p. injection of clozapine-n-oxide (CNO, Enzo Life Sciences) at a dosage of 0.3 mg/kg (working solution: 0.1 mg/mL in saline). On day 5, mice were perfused 45-55 minutes after the RI test. Behaviour was scored and analysed as previously explained.

#### Viral expression

ACC sections were checked along the rostro-caudal axis to verify the extent of (endogenous) viral expression. Three mice from the chemogenetics virus group had to be excluded due to virus expression in only one hemisphere (1 animal) and expression far beyond ACC into retrosplenial cortex (2 animals).

#### Immunohistochemistry

We assessed the number of active cells in ACC as well as downstream subcortical structures (amygdala, LH, VMH and MD) with a marker for cFos. For procedure please see the section on “Functional activation in ACC”.

