## Supplementary figures and images for "Anterior cingulate cortex hypofunction causes anti-social aggression in mice"

### Supplementary Material Figure 1

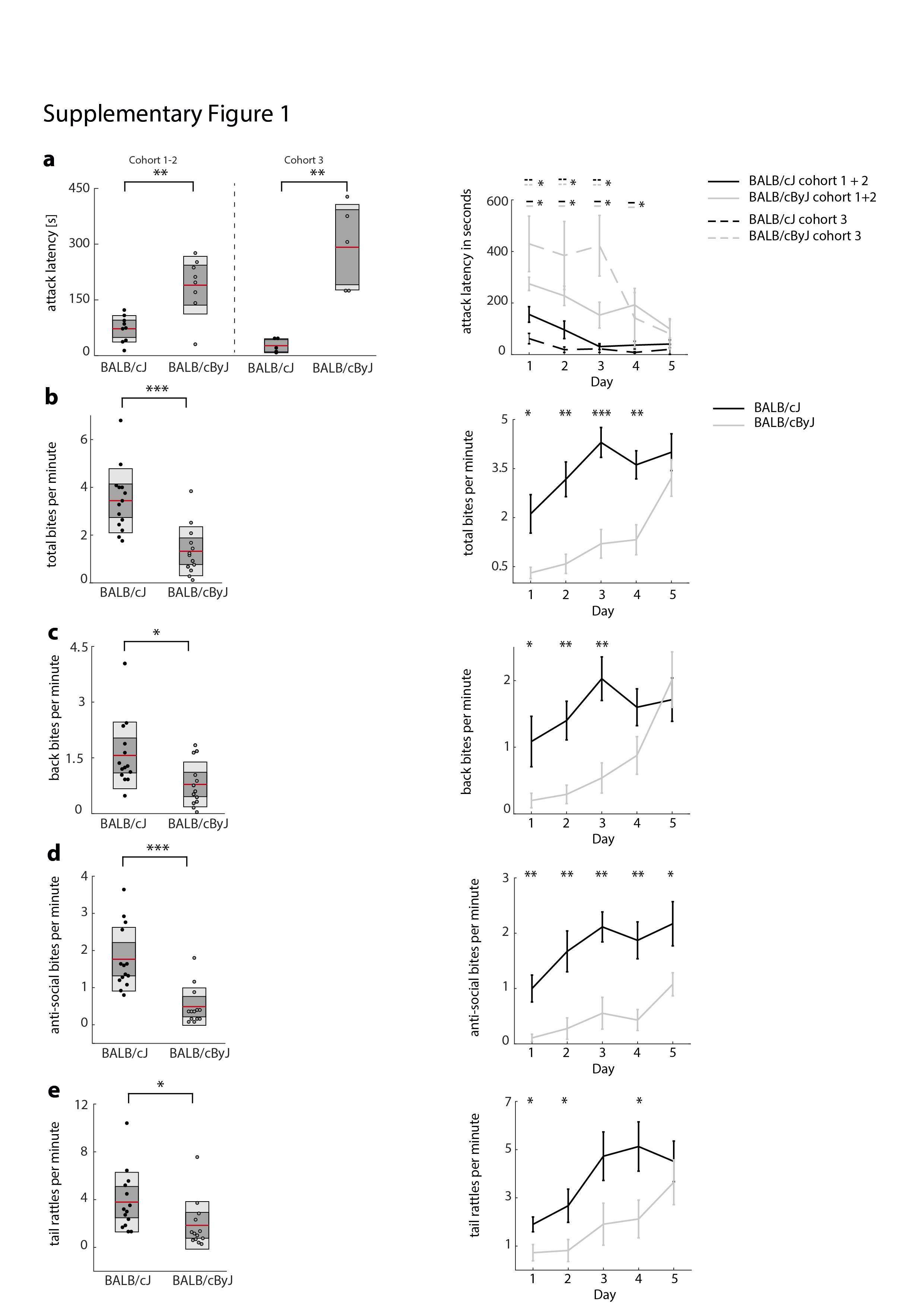

### Supplementary Material Figure 2

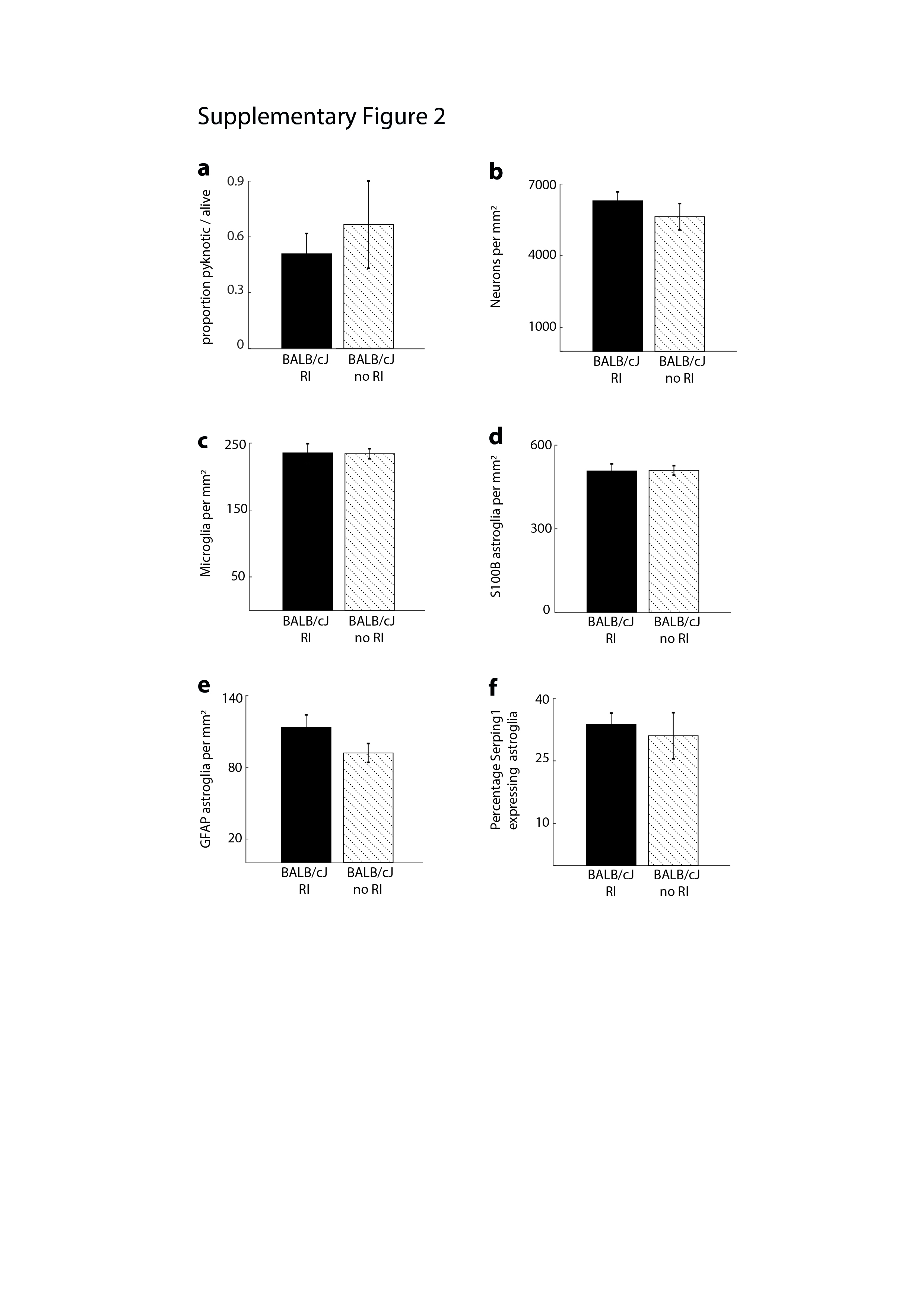

### Supplementary Material Figure 3

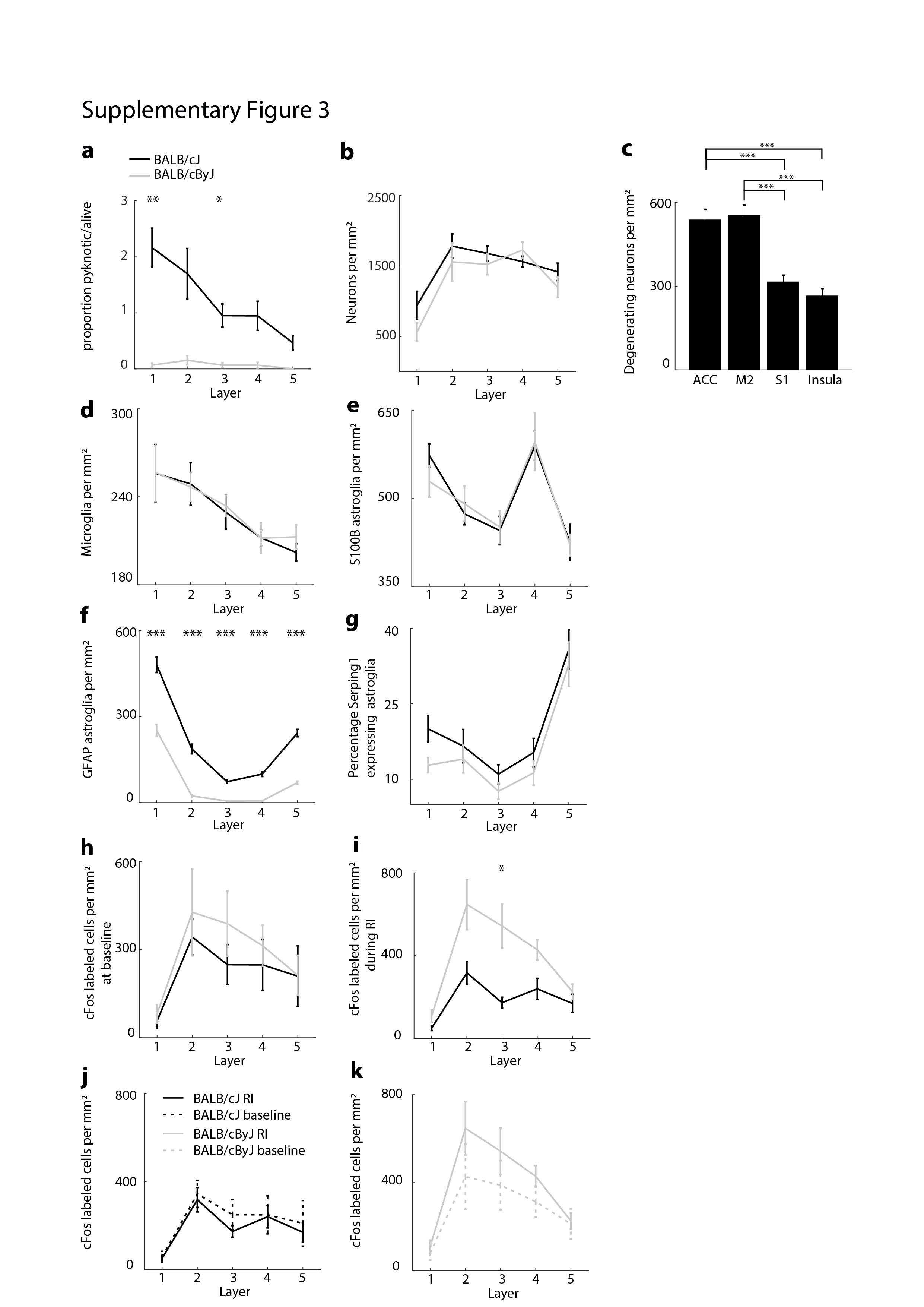

### Supplementary Material Figure 4

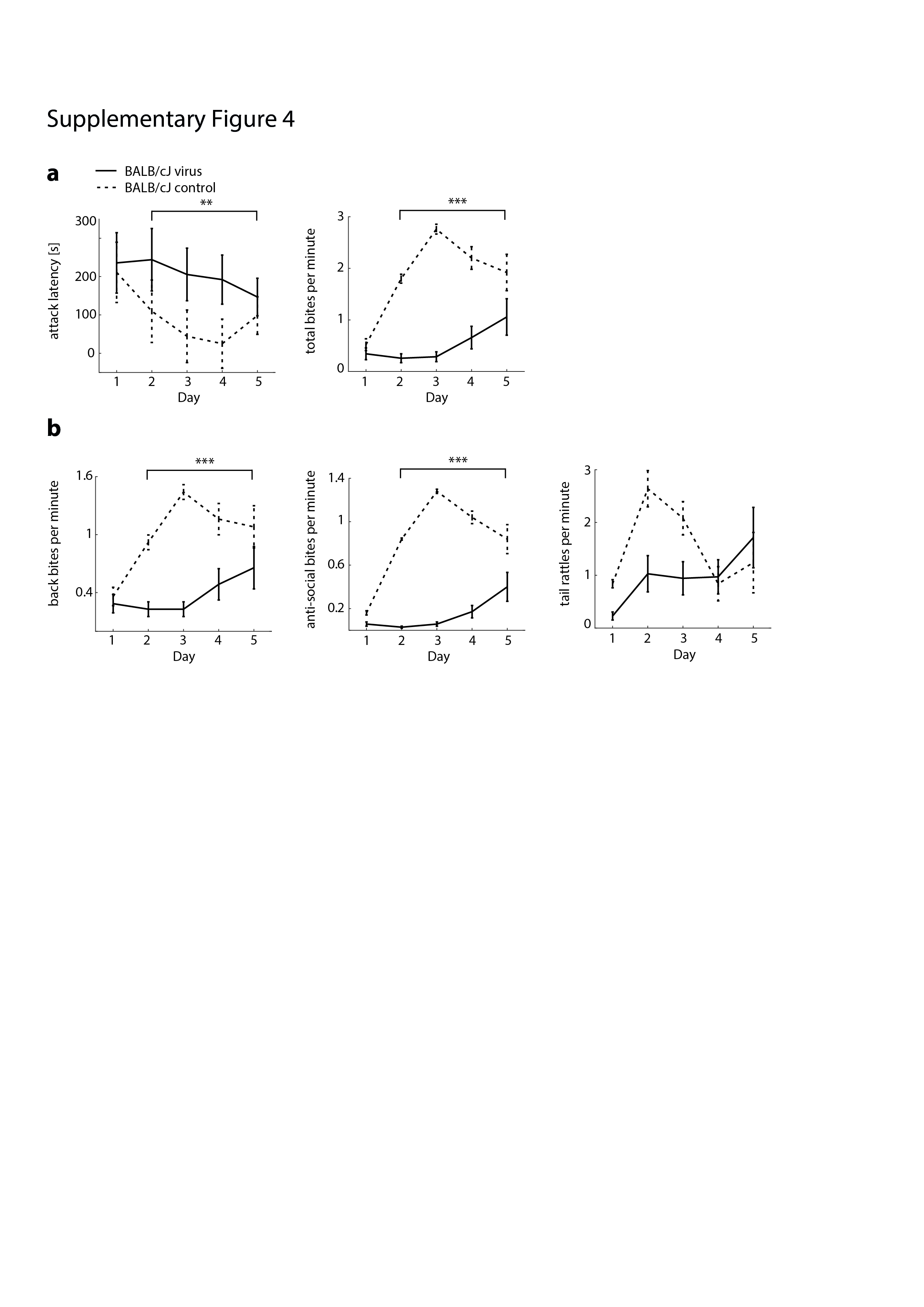

### Supplementary Material Figure 5

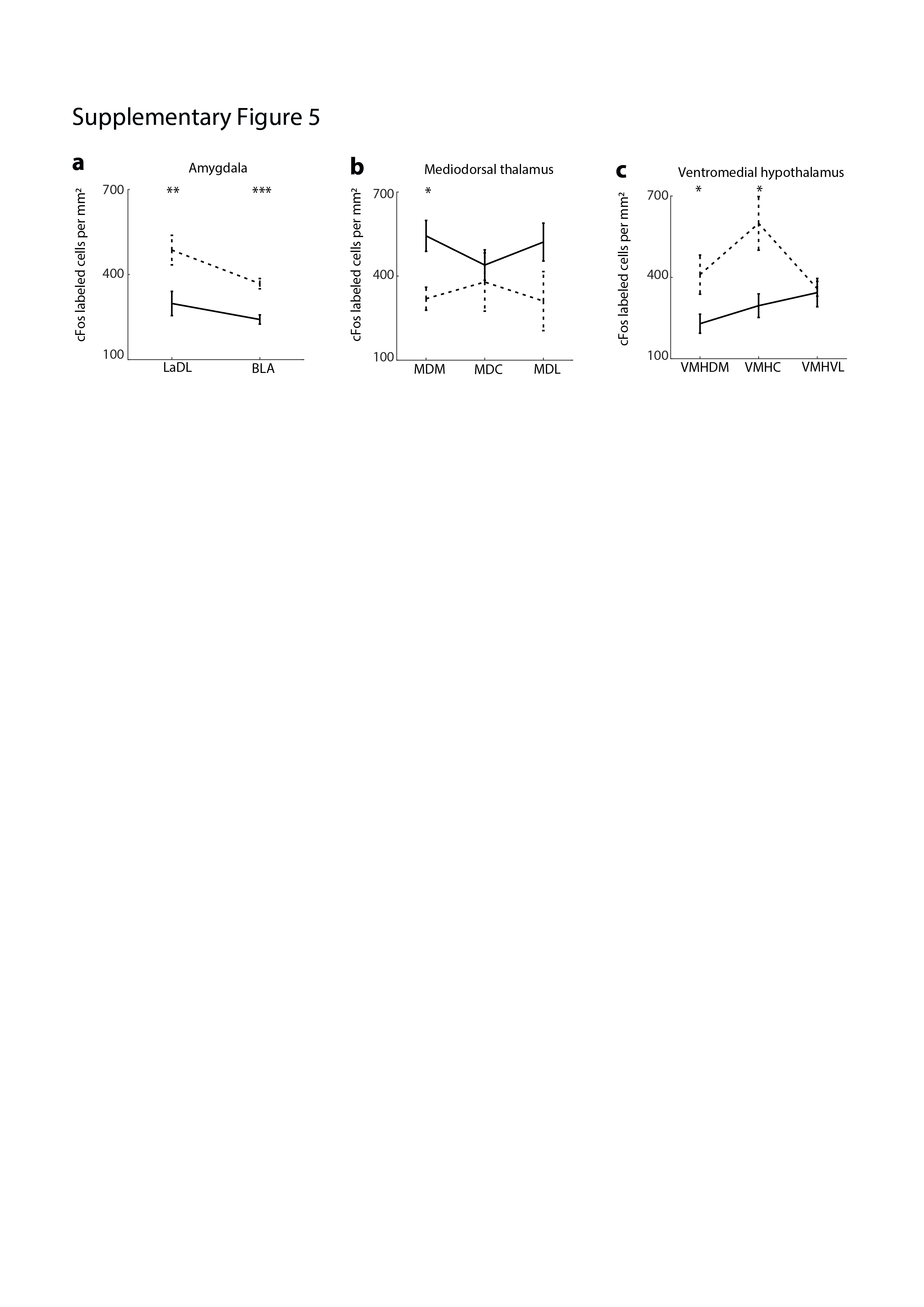

### Supplementary Material Figure 6

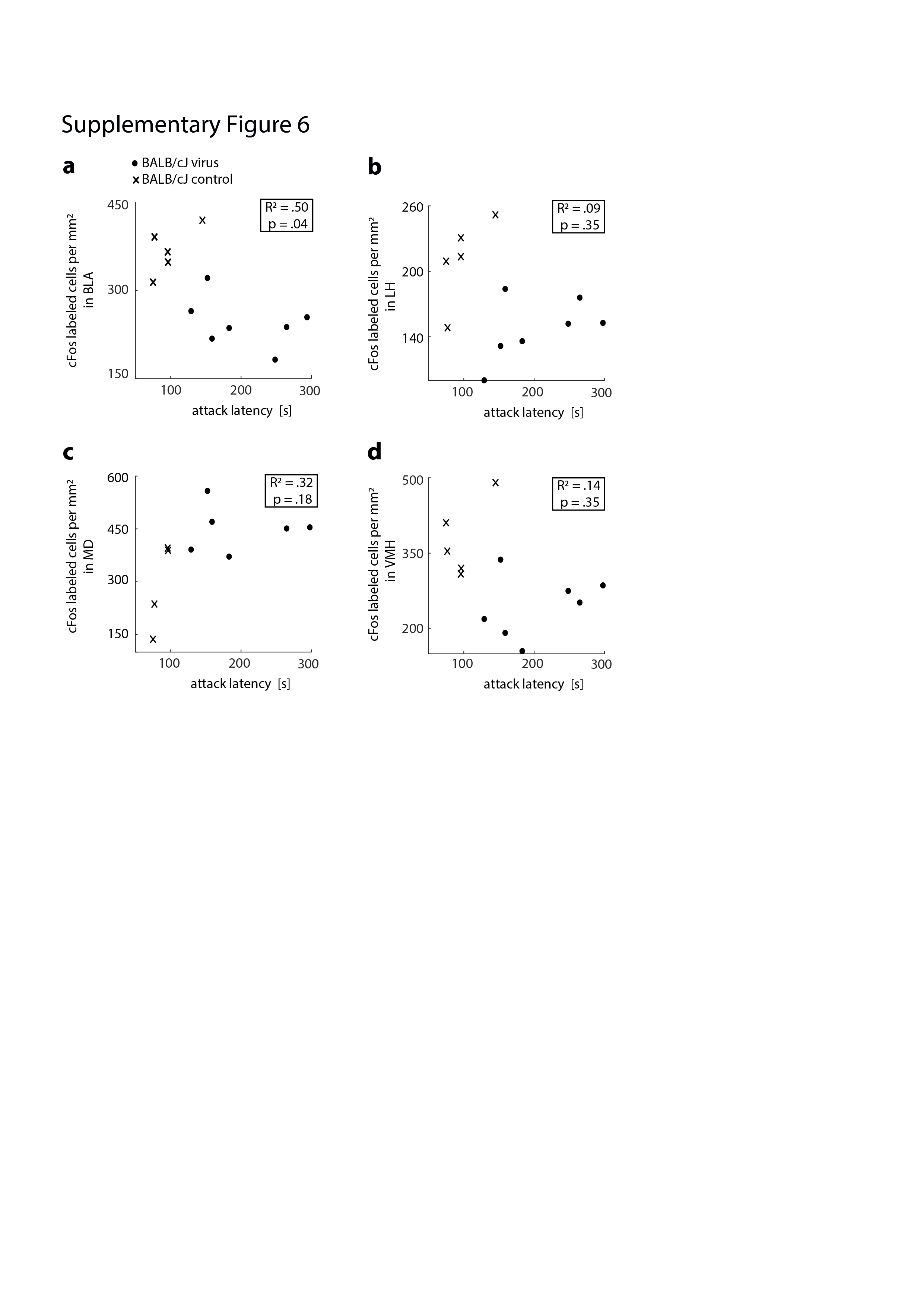
