## Supplementary Material for "Anterior cingulate cortex hypofunction causes anti-social aggression in mice"

**Supplementary material: Methods**

*Structural and functional measures of M2*. From each mouse the M2 sections were chosen at approximately the same anatomical level (AP 0.85). Sections were from the same mice as for ACC. M2 was defined according to [43] and we followed exactly the same approach as for the ACC sections. First, we determined the proportion of pyknotic / healthy neurons and then neuronal density. The same methodology and analysis was used as explained for ACC (section: Material & methods, Cytoarchitecture ACC). Second, we stained for glial markers, utilising the same antibodies as for ACC. At last, functional measures with cFos were obtained exactly as done for ACC.

*Fluorojade C staining*. The Fluoro-Jade C (FJC) Ready-to-Dilute Staining Kit for identifying Degenerating Neurons from Biosensis (Catalogue number: TR-100-FJ) was used to identify degenerating neurons in ACC, M2, S1 and insula. Sections were stained according to manufacturer's protocol with a 2-minute potassium permanganate interval and a 10-minute FJC incubation.

**Supplementary material: Figure legends**

*Figure 1*. Behavioural metrics of aggression. **a** Left: Average attack latencies shown across the cohorts. Right: Attack latency across all 5 days of the RI test. BALB/cJ and BALB/cByJ mice of cohort 3 had a maximum of 10 minutes to show a first attack, while mice of cohort 1 & 2 had a maximum of 5 minutes. This was done to ensure that control mice did not just need longer times to show aggressive behaviour. See in-figure legend. **b** Left: Average total bites. Black dots: BALB/cJ mice, grey dots: BALB/cByJ mice. Shown are average (red line), 95% confidence interval (dark-grey area) and 1 SD (light grey area). Right: Total bites across all days of the RI test. Black line: BALB/cJ mice, grey line: BALB/cByJ mice. Shown are average and SEM per day of the RI test. **c** Same as B for back bites. **d** Same as b for anti-social bites. **e** Same as b for tail rattles. * $p < .05$, ** $p < .01$, *** $p < .001$.

Figure 2. Structural measures RI vs baseline. **a** Proportion pyknotic neurons / healthy neurons in ACC of BALB/cJ mice that participated in the RI test vs those that did not (baseline). Shown are average and SEM, see in-figure legend. **b** Same as A for neuron density per $mm^2$. **c** Same as a for microglia per $mm^2$. **d** Same as a for S100B positive astroglia per $mm^2$. **e** Same as a for GFAP positive astroglia per $mm^2$. **f** Same as a for percentage toxic astroglia. * $p < .05$, ** $p < .01$, *** $p < .001$.

*Figure 3*. Structural and functional measures of control cortical areas. **a** Proportion pyknotic neurons / healthy neurons in secondary motor cortex (M2). Black line: BALB/cJ mice, grey line: BALB/cByJ mice. Shown are average and SEM per layer. **b** Same as a for neuron densities. **c** Degenerating neurons across ACC, M2, Somatosensory cortex (S1) and insula in BALB/cJ mice. Shown are average and SEM. See in-figure legend. **d** Same as a for microglia per $mm^2$. **e** Same as a for S100B positive astroglia per $mm^2$. **f** Same as a for GFAP positive astroglia per $mm^2$. **g** Same as a for percentage toxic astroglia. **h** The number of cFos labelled cells in per $mm^2$ at baseline. Black line: BALB/cJ mice, grey line: BALB/cByJ mice. Shown are average and SEM per layer. **i** Same as h for the number of cFos labelled cells per $mm^2$ during the last day of the RI test. **j** Comparison of activity at baseline vs RI for BALB/cJ mice. See in-figure legend. **k** Same as j for BALB/cByJ mice. * p < .05, ** p < .01, *** p < .001.

*Figure 4*. Behavioural effects of chemogenetic manipulation across R-I test days. **a** Left: Attack latency across the 5 days of the RI test. Black line: BALB/cJ mice injected with chemogenetics virus, dashed line: BALB/cJ mice injected with control virus. Shown are average and SEM per day of the RI test. Right: Same for total bites. **b** Same as a for back bites (left), anti-social bites (middle) and tail rattles (right). * p < .05, ** p < .01, *** p < .001.

*Figure 5*. cFos measures across sub-regions of ACC downstream areas. **a** Amygdala and its sub-regions. Black line: BALB/cJ mice injected with chemogenetic virus, dashed line: BALB/cJ mice injected with control virus. Shown are average and SEM. See in-figure legend. **b** Same as a for mediodorsal thalamus. **c** same as a for ventromedial hypothalamus. * p < .05, ** p < .01, *** p < .001.

Figure 6. Attack latency and activity in subcortical areas. **a** Correlation between BLA activity and average attack latency. Black dots: Mice injected with chemogenetic virus, crosses: mice injected with control virus. See in-figure legend. **b** Same as a for LH. **c** Same as a for MD. **d** Same as a for VMH.
